# Beyond IgG: Oral Secretory IgA as a new efficient modality for cancer immunotherapy

**DOI:** 10.64898/2026.11.23.768496

**Authors:** M. Brousse, A. Pintat, O. Favre-Bulle, B. Chanut, S. Paul, C. Genevois, C. Varon, B. Lejeune, JM. Herbert, A. Cuvillier

## Abstract

**BACKGROUD:** Monoclonal antibody therapies have revolutionized cancer treatment but are almost exclusively based on intravenously administered IgG antibodies. Here, we developed and evaluated the first orally delivered anti-CEACAM5 secretory IgA (SIgA) for colorectal cancer.

**METHODS:** The pharmacokinetics, biodistribution, antitumor efficacy and immune effects of oral anti-CEACAM5 SIgA (ABC-101) were evaluated in an orthotopic metastatic human colorectal cancer model. Its therapeutic activity was compared with cetuximab, as monotherapy and in combination with irinotecan- or FOLFIRI-based chemotherapy along with that of an anti-CEACAM5 IgG1 sharing the same antigen-binding domain. Tumor-associated FcαRI (CD89)-expressing myeloid populations were characterized at primary and metastatic sites. ABC-101 activity was evaluated on a second human cancer cell line, *in vivo* in an orthotopic human gastric cancer model.

**RESULTS:** Oral anti-CEACAM5 SIgA (ABC-101) exhibited a unique pharmacokinetic profile, selectively accumulating in CEACAM5-positive primary tumors and metastases while maintaining minimal systemic exposure. Compared with the anti-CEACAM5 IgG1 and cetuximab, oral SIgA achieved superior inhibition of primary tumor growth and metastatic dissemination. It also enhanced the efficacy of irinotecan- and FOLFIRI-based chemotherapy, outperforming the FOLFIRI/cetuximab standard-of-care comparator. Mechanistically, ABC-101 remodeled the tumor immune microenvironment by promoting the recruitment of FcαRI (CD89)-expressing neutrophils, monocytes, macrophages, and dendritic cells at both primary and metastatic sites, consistent with enhanced innate antitumor activity. Oral ABC-101 shows targeted activity against CEA^high^ MKN-45 cells, reducing markers of cell proliferation and migration.

**CONCLUSIONS:** These findings establish therapeutic SIgA as a first-in-class oral antibody immunotherapy combining selective tumor targeting, innate immune activation, and enhanced chemotherapy efficacy. More broadly, this work establishes mucosal immunity as a promising therapeutic frontier, demonstrating that orally delivered antibodies can enable systemic control of metastatic digestive cancers.

## WHAT IS ALREADY KNOWN ON THIS TOPIC

• Monoclonal antibody-based cancer immunotherapy relies almost exclusively on systemically administered IgG antibodies. While secretory IgA (SIgA) is known to be the predominant antibody isotype at mucosal surfaces engaging FcαRI/CD89-expressing myeloid effector cells, its immunotherapeutic potential as an oral chemotherapy remains largely unexplored.

## WHAT THIS STUDY ADDS

• This study provides the first preclinical evidence that an orally administered tumor targeting SIgA can control both primary and metastatic colorectal cancer. Oral ABC-101 activity was confirmed on a second human digestive cancer model. Anti-CEACAM5 SIgA (ABC-101) combines tumor-selective tissue accumulation and prolonged retention with recruitment of CD89-expressing myeloid effector cells, achieving greater antitumor activity than IgG antibodies while enhancing chemotherapy efficacy.

## HOW THIS STUDY MIGHT AFFECT RESEARCH, PRACTICE OR POLICY

• These findings introduce oral SIgA as a new antibody immunotherapy modality and support the intestinal mucosa as a potential gateway for the treatment of local and metastatic cancer. They provide a rationale for the clinical development of orally delivered SIgA treatments and for extending this approach to other tumor-associated targets and mucosal cancers.

## INTRODUCTION

Colorectal cancer (CRC) remains one of the leading causes of cancer-related mortality worldwide, with nearly two million new cases diagnosed annually and a rising incidence among adolescents and young adults (1). Despite major advances in surgery, chemotherapy and targeted therapies, metastatic CRC remains associated with poor outcomes. Immune checkpoint inhibitors have transformed the treatment of mismatch repair-deficient/microsatellite instability-high (dMMR/MSI-H) tumors but provide limited benefit in microsatellite-stable (MSS) disease, which accounts for approximately 85% of CRC cases. New immunotherapeutic strategies for advanced CRC therefore remain a major unmet clinical need. Antibody-based cancer immunotherapy has been built almost exclusively around IgG. Although monoclonal IgG antibodies have transformed cancer treatment, their pharmacology is intrinsically systemic: prolonged circulation can contribute to off-target toxicity, while penetration across epithelial barriers and into solid tumors remains challenging. In contrast, the therapeutic potential of IgA — the predominant antibody class at mucosal surfaces — remains largely unexplored in oncology, despite mucosal tissues giving rise to nearly two-thirds of human cancers.

IgA possesses biological and pharmacological properties fundamentally distinct from IgG. It is the most abundantly produced antibody isotype in humans (approximately 66 mg/kg/day), the second most abundant in serum and the dominant immunoglobulin at mucosal surfaces. IgA exists as monomers in blood, dimers/polymers (dIgA/pIgA) within mucosal tissues and secretory IgA (SIgA) in mucosal lumen. Mucosal B cells predominantly produce J-chain-containing dIgA, which binds the polymeric immunoglobulin receptor (pIgR) and undergoes epithelial transcytosis. Cleavage of pIgR during apical release generates the secretory component, which remains associated with IgA to form SIgA (2,3). SIgA can also undergo retrograde transport across mucosal barriers through M-cell- and receptor-associated pathways involving Dectin-1 (4), SIGN-R3, SIGLEC-5 and epithelial CD71, facilitating delivery of immune complexes to dendritic cells through DC-SIGN (5). Hepatobiliary pIgR-mediated transport provides an additional route connecting systemic and gastrointestinal IgA compartments (6). Thus, unlike conventional IgG, SIgA is naturally adapted to trafficking, persistence, and immune surveillance across mucosal environments.

Importantly, IgA is not merely a transport or barrier antibody. It can support complement-dependent cytotoxicity, antibody-dependent cellular cytotoxicity (ADCC), phagocytosis (ADCP), and trogocytosis (ADCT). Several studies have shown that IgA antibodies can outperform their IgG counterparts against HER2-, EGFR- and CD20-expressing tumor cells, particularly with neutrophils as effector cells. This enhanced cytotoxic activity is largely mediated by FcαRI (CD89), whose engagement triggers potent ITAM-dependent signaling (7). IgA therefore provides access to a powerful FcαRI-positive myeloid effector compartment poorly exploited by conventional IgG therapeutics.

Emerging evidence from human cancer further challenges the IgG-centric view of antitumor humoral immunity. Tumor-associated tertiary lymphoid structures have recently been shown to harbor highly oligoclonal; affinity-matured IgA responses and a recombinant dimeric IgA cloned directly from B cells infiltrating a human tumor suppressed growth of the corresponding autologous tumor *in vivo*. These observations establish IgA as a naturally occurring component of human antitumor immunity and are consistent with evidence that dIgA can undergo pIgR-dependent transcytosis through tumor cells and induce through their SIgA form biological changes detrimental to tumor survival (8).

SIgA may also bridge innate and adaptive antitumor immunity. Beyond FcαRI-dependent myeloid cytotoxicity, SIgA interacts with immunoregulatory receptors. FCRL3, expressed by subsets of B lymphocytes and FOXP3⁺ regulatory T (Treg) cells, has been reported to interact with SIgA, potentially reducing Treg-mediated suppression and restoring CD4⁺ and CD8⁺ effector T-cell proliferation (9). IgA isoforms can also modulate dendritic-cell activation and promote cytotoxic CD8⁺ T-cell responses. Furthermore, pIgR-expressing ovarian cancer cells can internalize tumor-associated dIgA and locally generate SIgA, demonstrating that the IgA/SIgA pathway can operate directly within the human tumor microenvironment. SIgA may therefore combine tumor targeting, FcαRI-dependent myeloid cytotoxicity and modulation of adaptive immunity within a single antibody architecture.These properties are particularly attractive for targeting carcinoembryonic antigen (CEA/CEACAM5). CEA is expressed at low levels on normal gastrointestinal epithelial cells but is strongly overexpressed on the apical surface of many adenocarcinomas (10). In advanced colorectal cancer (11,12), CEA is detected in approximately 98% of tumors and is associated with disease progression, poor prognosis, and recurrence (13,14). Its high prevalence in malignant tissue and comparatively restricted expression in healthy tissues have made CEA an attractive therapeutic target for antibodies, antibody-drug conjugates, CAR-T cells and vaccines. Yet, despite decades of therapeutic development, secretory IgA has never been exploited to target CEA-positive malignancies.

Here, we address this question using ABC-101, a tumor-specific anti-CEA secretory IgA administered orally, in an orthotopic metastatic human colorectal cancer model. ABC-101 remains biologically active following oral administration, displays a pharmacokinetic profile fundamentally distinct from intravenous IgG, and selectively accumulates and persists, despite limited systemic exposure. Oral ABC-101 suppresses primary tumor growth and metastatic dissemination as monotherapy, with greater efficacy than intravenous cetuximab and a paratope-matched IgG1 counterpart while further enhancing therapeutic activity of chemotherapy. Its antitumor activity is associated with a marked recruitment of FcαRI/CD89-positive myeloid effector cells into both primary and metastatic lesions, which was not observed following treatment with the standard-of-care immunotherapy cetuximab, an anti-EGFR IgG.

Together, these findings provide, to our knowledge, the first *in vivo* evidence that an orally administered secretory IgA can operate as an effective anticancer immunotherapy. Beyond ABC-101 and CEA-positive digestive cancers, they introduce oral SIgA as a distinct therapeutic modality combining mucosal delivery, tumor-selective tissue retention and IgA-specific immune effector functions, opening a previously unexplored field in antibody-based cancer immunotherapy.

## KEY MESSAGE

ABC-101 introduces a new paradigm in cancer immunotherapy. We demonstrate that an orally administered secretory IgA can selectively target primary and metastatic tumors while triggering mucosal immunity. This work positions secretory IgA as a new generation of therapeutic antibodies with the potential to overcome the current limitations of conventional IgG antibodies redefining immunotherapy for the treatment of metastatic solid tumors.

## RESULTS

### Generation of ABC-101, a monoclonal CEACAM5-specific humanized Secretory IgA

To generate a monoclonal antibody specific for carcinoembryonic antigen (CEA/CEACAM5), HAMIGA™ transgenic mice were immunized with full-length recombinant human CEA. Following spleen isolation, approximately 8.5 × 10⁷ splenocytes were recovered and fused with myeloma cells, yielding approximately 600 hybridoma clones. Hybridoma supernatants were initially screened by CEA-specific ELISA and subsequently evaluated by cell-based ELISA using fixed human colorectal cancer cells expressing CEA. The hybridoma clone exhibiting the highest CEA-binding activity was selected for molecular characterization.

The variable heavy (VH) and light (VL) domains of the selected antibody were sequenced and humanized by grafting the complementarity-determining regions (CDRs) onto human germline framework sequences. The humanized VH sequence was cloned upstream of the human IgA1 constant region, whereas the VL sequence was cloned upstream of the human κ light-chain constant region.

The purified SIgA1 antibody, designated ABC-101, was analyzed by SDS–NuPAGE under both non-reducing and reducing conditions (Figure 1a). Under non-reducing conditions, a predominant band of approximately 350 kDa accounted for 93% of the total protein, consistent with the expected molecular weight of intact SIgA1. Under reducing conditions, four protein bands were detected, corresponding to the J chain (20–25 kDa; one N-glycosylation site), the κ light chain (25–28 kDa), the IgA1 heavy chain (55–60 kDa; five to eight O-glycosylation sites and two N-glycosylation sites), and the secretory component (75–80 kDa; seven N-glycosylation sites).

**Figure 1:**
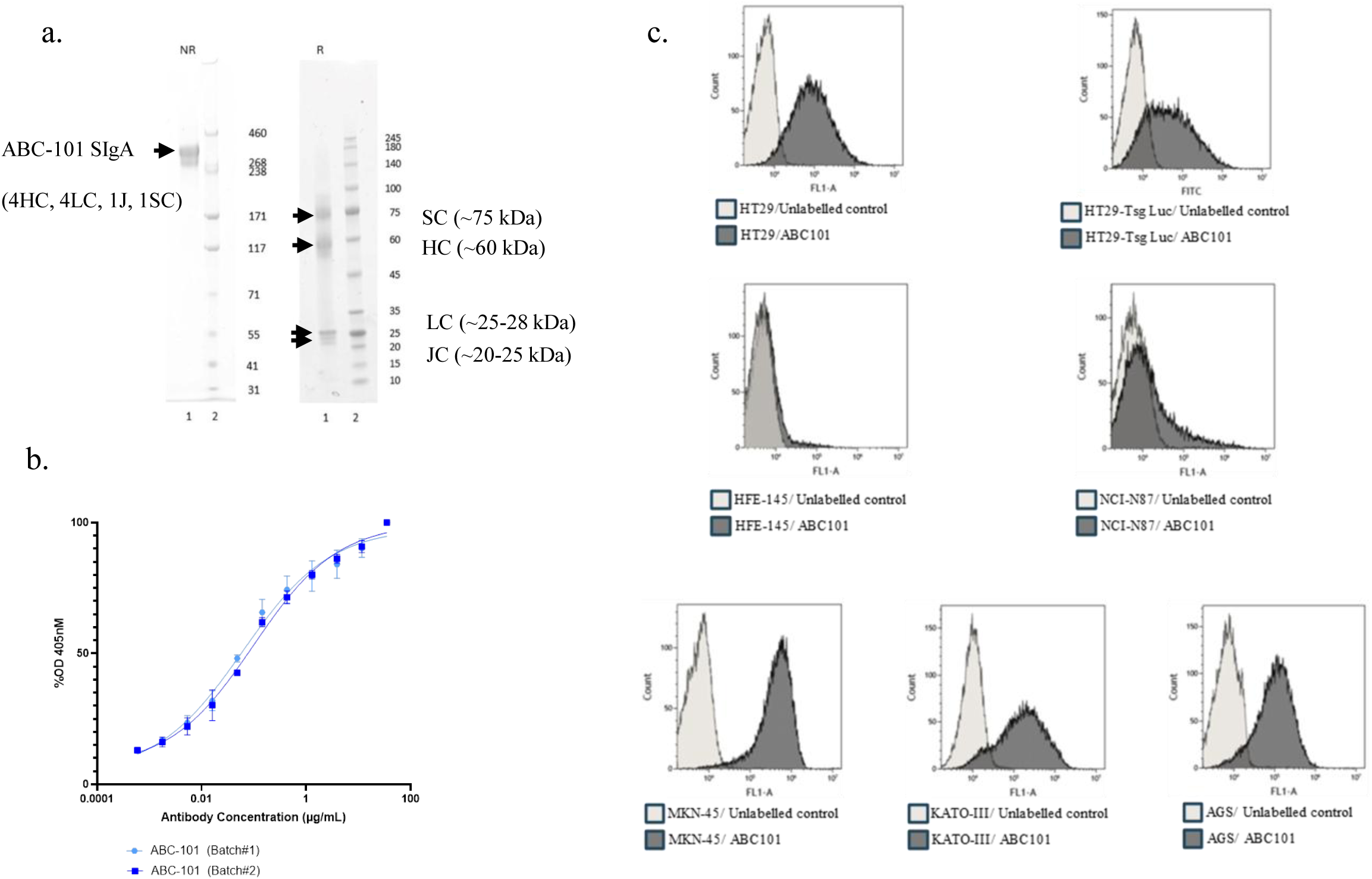
Anti-CEACAM-5 specificity of ABC-101. a. sIgA and sub-units separation by Nu-Page gel, Analysis of purified ABC-101 Batch #1 by Nu-PAGE Coomassie staining, 2µg of protein was loaded onto a gel and analyzed under non-reducing (NR) or reducing conditions, b. ELISA evaluation of hCEACAM-5 avidity of two production batches of ABC-101, c. Cell cytometry validation of CEACAM-5 recognition by ABC-101 on CRC tumor cell lines (HT-29) and gastric tumor cells (HFE-145, NCI-N87, MKN-45, KATO-III, AGS)

The binding avidity of ABC-101 for native CEA purified from human colorectal adenocarcinoma extracts was determined by ELISA (Figure 1b). Two independent production batches manufactured two months apart exhibited identical binding characteristics, with EC₅₀ values of 0.03 ± 0.01 µg/mL for Batch #1 and 0.03 ± 0.01 µg/mL for Batch #2. .

### ABC-101 selectively targets tumor cells with moderate-to-high CEACAM5 expression

ABC-101 bound strongly to HT-29, HT-29_Luc, AGS, MKN-45, and KATO-III cells, cell lines expressing moderate to high levels of membrane-associated CEACAM5 whereas no detectable binding was observed in HFE-145 or NCI-N87 cells, expressing very low levels of the antigen (Figure 1c.).

### ABC-101 remains highly stable under physiological and GastroIntestinal conditions

The stability of ABC-101 under physiological conditions was evaluated using ^125^I-labeled ABC-101 incubated in fresh mouse blood at 37°C (Figure 2a.). After 1 hour, 98% of the radiolabeled antibody remained intact (83.3 ± 0.6% at T₀ versus 81.8 ± 0.1%). After 24 hours, 79.7 ± 0.3% of total radioactivity remained associated with the high-molecular-weight ABC-101 peak, compared with 83.3 ± 0.6% at T₀, corresponding to preservation of approximately 96% of the initially radiolabeled protein.

**Figure 2:**
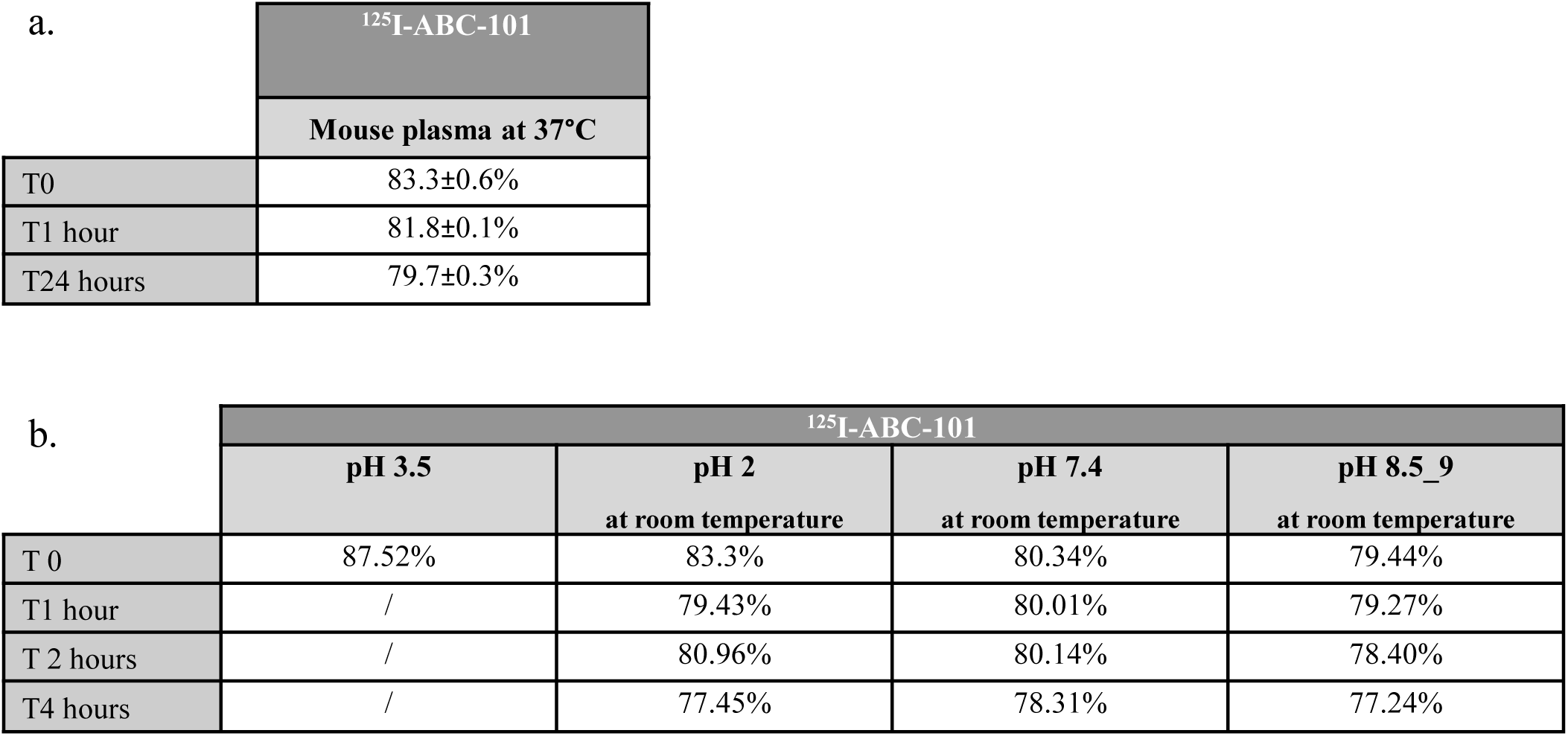
Stability test of ABC-101. a. Table 1: Radiochemical purity of ^125^I-ABC101 in mouse plasma at 37°C Data expressed as percentage of Iodine-125 still associated to protein (Chelatec source). Plasma stability of ^125^I-ABC-101 was assessed using mouse plasma incubated at 37°C. b. Table 2: Radiochemical purity of ^125^I-ABC-101 in three pH-stabilized buffer at RT – Data expressed as percentage of Iodine-125 still associated to protein (ITLC analyses) Buffer stability of ^125^I-ABC-101 as labelling solution was assessed using three pH-stabilized buffers at room temperature. The concentration of ^125^I-ABC-101 in buffer corresponded to a 1/10 dilution of the radiolabelled product in buffers

ABC-101 stability was also evaluated under pH conditions representative of the gastrointestinal tract. ^125^I-ABC-101 was incubated for up to 4 hours at acidic (pH 2.0), neutral (pH 7.4), or alkaline (pH 8.5–9.0) conditions, corresponding to the gastric, duodenal/colonic, and jejunal/ileal environments, respectively. At pH 2.0, the ABC-101 peak represented 77.45% of total radioactivity after 4 hours versus 83.3% at T₀, corresponding to approximately 93% preservation of the initial radiolabeled antibody. Its retention time (17.37 min) remained comparable to the reference material (17.16 min), supporting preservation of the molecular species (Figure 2b.). Under neutral and alkaline conditions, 97.5% and 97.2%, respectively, of radiolabeled ABC-101 remained intact after 4 hours. Overall, these results demonstrate that ABC-101 remains highly stable in blood and across the physiological pH range encountered in the gastrointestinal tract.

### Oral ABC-101 combines limited systemic exposure with selective and prolonged tumor retention

The pharmacokinetic and biodistribution profiles of orally administered ABC-101 were evaluated in healthy SCID mice and compared with ABC-102, a human IgG1 carrying the same antigen-binding variable domains and identical affinity for human CEACAM5. Both antibodies were radiolabeled with ^125^I and administered at identical doses, orally for the SIgA ABC-101 and intravenously for ABC-102, its IgG counterpart .

Following oral administration, ^125^I-ABC-101 reached a Tmax of approximately 4 hours, with an elimination half–life of 6–8 hours and declined to near-background circulating levels within 24 hours. In contrast, IV ^125^I-ABC-102 remained detectable throughout 96 hours, with circulating levels decreasing by approximately one order of magnitude (Figure 3a).

**Figure 3.**
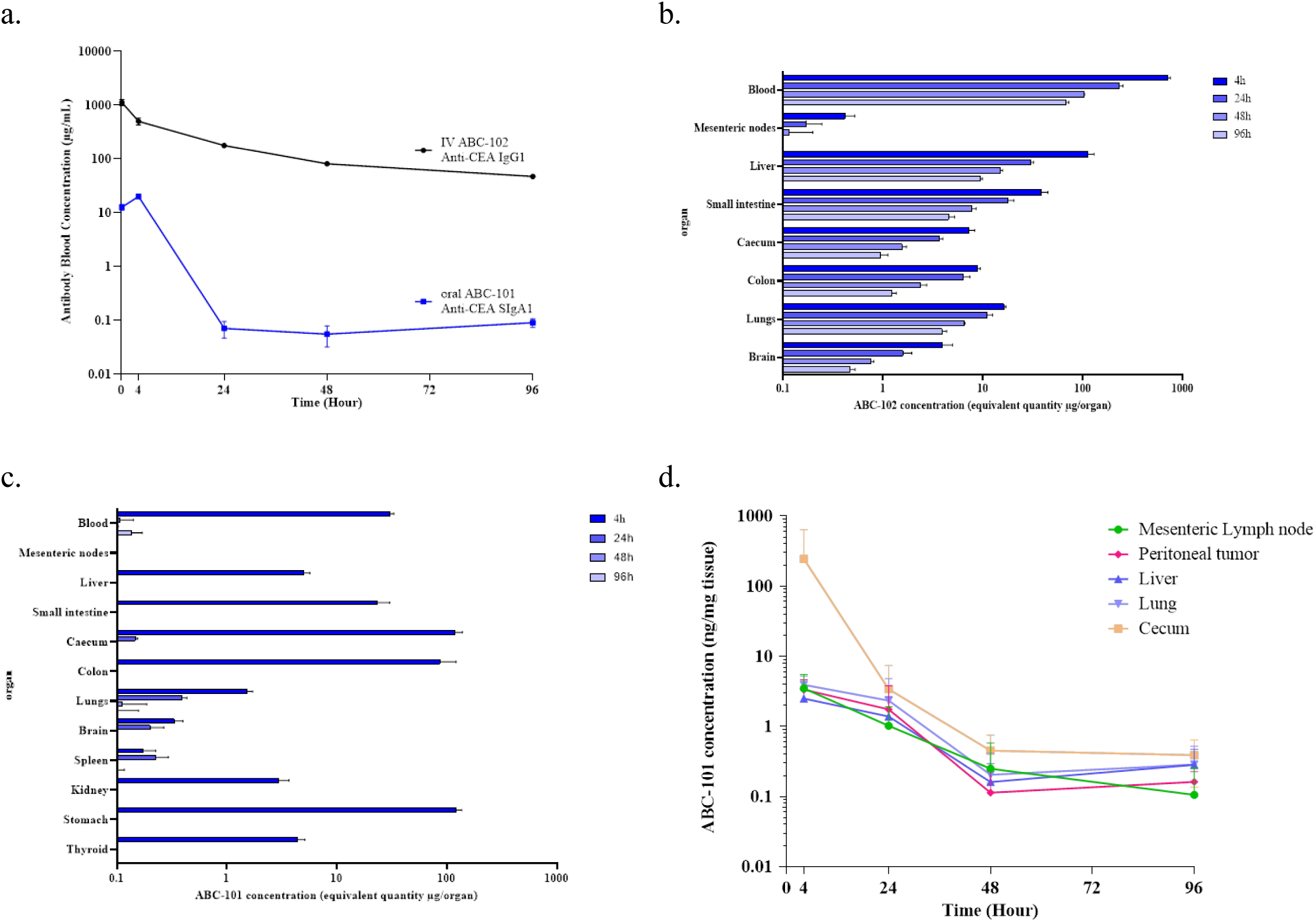
Pharmacokinetics and biodistribution of oral ABC-101 in healthy and tumour-bearing mice, and comparison with the tissue distribution of ABC-102, an epitope-matched IgG. a. Blood pharmacokinetics of ABC-101 and ABC-102 in healthy mice, b. ABC-101 tissue distribution in healthy mice, c. ABC-102 tissue distribution in healthy mice, d. Tissue distribution of ABC-101 in tumor bearing mice

In healthy mice, ABC-101 was predominantly localized in the gastrointestinal tract. At 4 hours, 122 ± 14 µg was detected in the stomach, 120 ± 18 µg in the cecum, 86 ± 35 µg in the colon, and 23 ± 7 µg in the small intestine, representing approximately 40% of the administered dose (Figure 3b). Much lower levels were detected in the liver (5.03 ± 0.65 µg), kidneys (2.97 ± 0.69 µg), lungs (1.52 ± 0.20 µg), and spleen (0.17 ± 0.05 µg). At 24 hours, only trace amounts of SIgA remained detected in the lungs, cecum, brain, and spleen, with no substantial signal thereafter. In contrast, intravenous administration of IgG ABC-102 resulted in sustained systemic exposure, with detectable antibody levels persisting in all vascularized organs throughout the 96 hours (Figure 3c).

ABC-101 distribution was also evaluated in SCID mice bearing orthotopic HT-29_Luc colorectal tumors and metastases in the peritoneum, mesenteric lymph nodes, liver, and lungs. In primary cecal tumors, ABC-101 peaked at approximately 4 hours at 243.6 ± 394.1 ng/mg and remained detectable at 96 hours at 0.39 ± 0.25 ng/mg (Figure 3d).

Importantly, ABC-101 also showed prolonged retention in metastatic tissues. At 24 hours, liver concentrations and lung concentrations were respectively 1.40 ± 1.27 ng/mg and 2.32 ± 2.18 ng/mg versus 0.138 ± 0.040 ng/mg and 0.12 ± 0.01 ng/mg in healthy mice tissues. Mesenteric lymph nodes showed approximately four orders of magnitude higher retention than healthy lymph nodes. Peritoneal metastases contained 3.34 ± 0.90 ng/mg at 4 hours, with ABC-101 remaining detectable up to 96 hours at 0.162 ± 0.005 ng/mg.

### ABC-101 strongly recognizes native CEA across human GastroIntestinal tumors

The ability of ABC-101 to recognize membrane-associated CEA was evaluated by immunohistochemistry using formalin-fixed, paraffin-embedded human tumor specimens obtained from patients with stage III rectal, esophageal, gastric adenocarcinoma, and stage III and IV colon adenocarcinoma. The staining pattern and intensity obtained with ABC-101 were compared with those produced by a routinely used diagnostic anti-CEA antibody on the Leica Bond RX automated immunostaining platform. Across all tumor types examined, ABC-101 generated a staining pattern comparable to that of the reference diagnostic antibody while consistently producing slightly stronger staining intensity (Figure 4a), indicating efficient recognition of native CEA in human tumor tissues.

**Figure 4:**
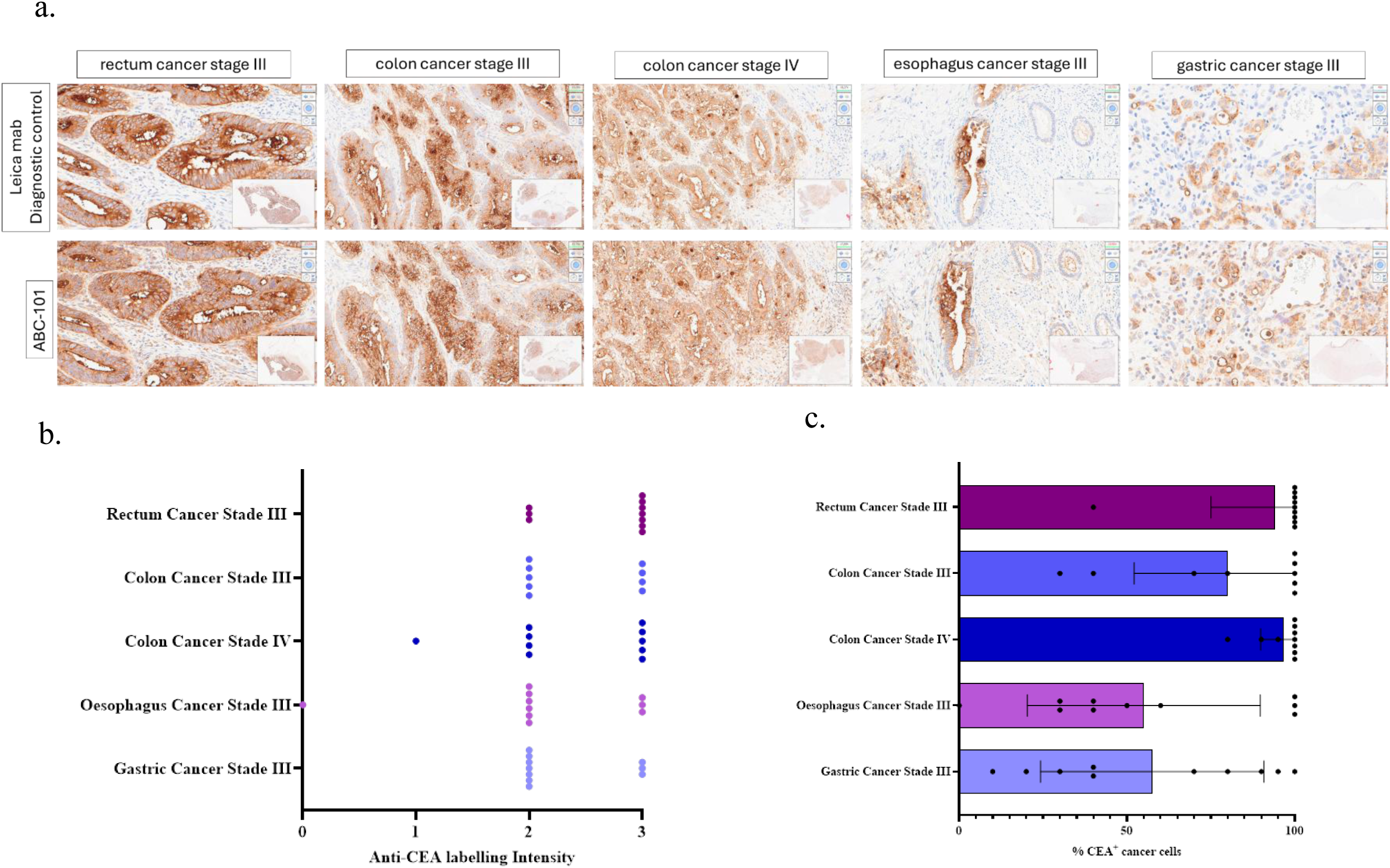
CEACAM-5 expression in human digestive cancers. a. Immuno-histochemistry validation of human CEACAM-5 labeling by ABC-101 on human cancer tissues from rectum cancer, colon cancer stage III, colon cancer stage IV, esophagus cancer and gastric cancer in comparison to labeling pattern of diagnostic validated anti-CEACAM-5 antibody, b. Percentage ofh CEACAM-5 positive cancer cells in human tumoral tissue, c. Intensity of hCEACAM-5 labeling in human tumoral tissue.

All positive colorectal specimens exhibited strong staining (up to a maximum intensity score of 3). In colon adenocarcinoma, staining intensity increased with tumor stage, with the median staining score rising from 2 in stage III tumors to 2.5 in stage IV tumors (Figure 4b). Positive CEA staining was observed in 94 ± 6% of stage III rectal adenocarcinomas, 80 ± 28% of stage III colon adenocarcinomas, and 96 ± 7% of stage IV colon adenocarcinomas. CEA expression was more heterogeneous in upper gastrointestinal malignancies (Figure 4c). Positive staining was detected in 55 ± 35% of tumor cells in stage III esophageal adenocarcinomas and in 57 ± 33% of tumor cells in stage III gastric adenocarcinomas. Among CEA-positive tumors, moderate staining predominated, whereas strong staining (intensity score 3) was observed in approximately 30–37% of esophageal and gastric adenocarcinomas.

### *In Vivo* transcytosis of orally administered ABC-101 through Peyer’s Patches

To investigate the intestinal transcytosis of orally administered ABC-101, a ligated ileal loop model containing a Peyer’s patch was used. Under anesthesia, ABC-101 was injected into the intestinal loop lumen, and the ileal segment was collected 1.5 hours later for immunohistochemical analysis using a PE-conjugated anti-human IgA antibody. ABC-101 was detected within the intestinal lumen, where part of the antibody remained associated with the mucus layer. In addition, ABC-101 was observed within the intestinal epithelium, throughout the lamina propria, and in the submucosal compartment. Positive staining was also detected in submucosal blood vessels, demonstrating that ABC-101 had crossed the intestinal epithelial barrier and reached the underlying vascular compartment (Figure 5, red staining, panel c.). These findings provide direct evidence that orally administered ABC-101 undergoes transcytosis across the intestinal mucosa *via* Peyer’s patches, thereby gaining access to the systemic circulation.

**Figure 5:**
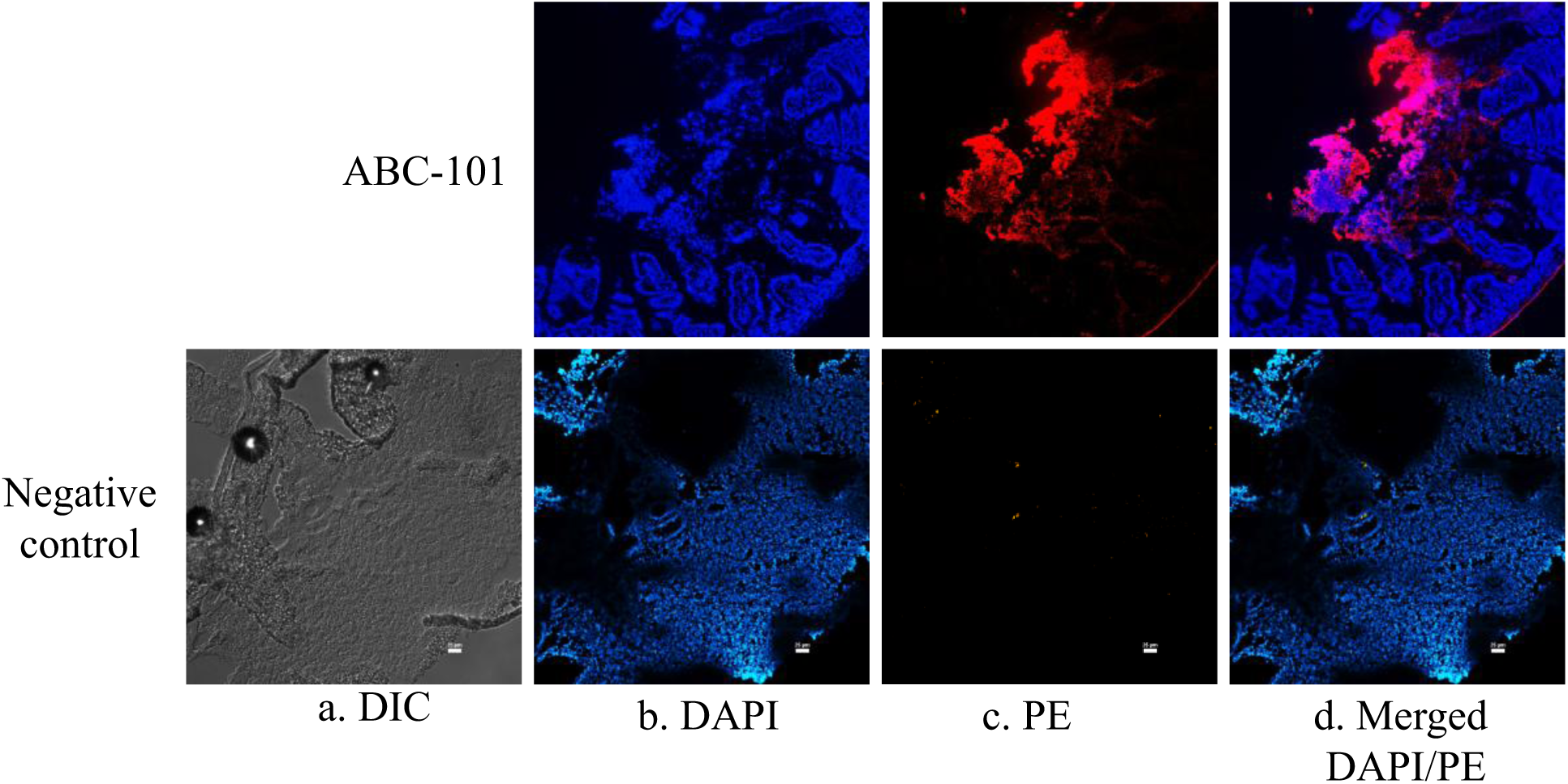
Reverse transcytosis of ABC-101 through intestinal Peyer patches. Representative microscopy images of Immunofluorescence detection of PE-labeled ABC-101 in ligated Peyer patch tissue 2h after intra-lumen injection. Panel a. DIC image acquired with a 10x magnitude objective. Nuclei are visualized by DAPI labeling (panel b.). IgA are visualized by anti-hIgA-PE labeled secondary antibody (panel c.). Merged image is shown in panel d.

### Orally delivered ABC-101 suppresses colorectal tumor growth more effectively than intravenous IgG therapies

The therapeutic efficacy of ABC-101 was evaluated in an orthotopic HT-29_Luc human colorectal cancer model established in transgenic Tg-hCD89 mice. Treatment was initiated when primary cecal tumors reached a bioluminescence signal of 5×10^6^ to 5×10^7^ photons/cm², corresponding to locally advanced tumors associated with metastases in mesenteric lymph nodes, liver, and lungs. At the experimental endpoint, extensive tumor infiltration precluded the isolation of discrete tumor masses. Therefore, the entire cecum (after removal of luminal contents), liver, and lungs were collected and weighed consistently across all groups for analysis. ABC-101 was administered orally by gavage at doses of 0.2 or 0.8 mg once daily, 5 days per week for 3 weeks. ABC-102, the human IgG1 counterpart of ABC-101 sharing identical antigen-binding variable domains, and cetuximab (Erbitux®, human IgG1 anti-EGFR) were administered intravenously at 4 mg the first week, followed by a 2,5mg dose once weekly for 2 weeks. Both oral doses of ABC-101 significantly reduced primary cecal tumor burden, with comparable efficacy observed at 0.2 mg and 0.8 mg. Compared with untreated controls, treatment with 0.8mg of ABC-101 allowed a decrease of mean tumor weight to 0.167 ± 0.035 g, showing a 59% drop. (0.404 ± 0.212 g; P < 0.0001) (Figure 6a.). Remarkably, oral ABC-101 outperformed both intravenous comparator antibodies; with primary tumor weight significantly lower in mice treated with ABC-101 than in those receiving intravenous cetuximab (0.167 ± 0.035 g *vs.* 0.241 ± 0.047 g; p = 0.0004) or ABC-102 (0.242 ± 0.048 g), corresponding to an additional reduction of approximately 31% relative to either IgG-based treatment. These findings demonstrate that orally administered ABC-101 effectively suppresses primary colorectal tumor growth and provides superior antitumor activity compared with intravenously administered IgG antibodies targeting either CEA or EGFR.

**Figure 6:**
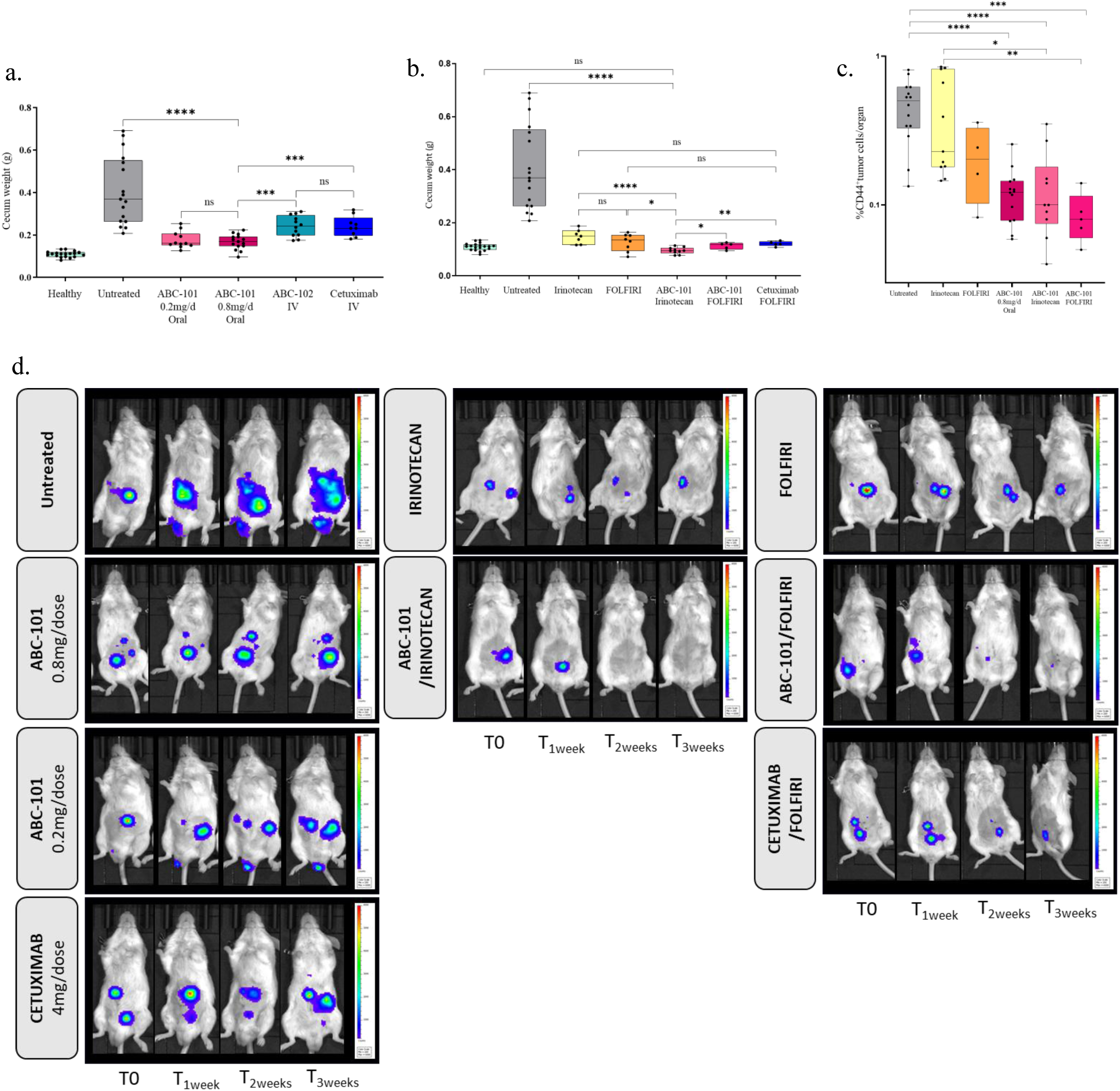
Therapeutic efficacy of oral ABC-101 treatment. a. Monotherapy efficacy on primary tumor weight after oral administration of ABC-101 0.2 or 0.8mg/dose; ABC-102 (ABC-101 counterpart anti-CEA IgG1) 4mg/dose given IV or Cetuximab 4mg/dose given IV, b. Therapeutic combinations efficacy on primary tumor weight after oral administration of ABC-101 0.2 or 0.8mg/dose; ABC-102 (ABC-101 counterpart anti-CEA IgG1) 4mg/dose given IV or Cetuximab 4mg/dose given IV, c. Percentage of CD44^+^-HT29/organ in pulmonary metastases, after oral monotherapy with ABC-101 0,8mg/dose, or combination therapy with FOLFIRI or IRINOTECAN, d. Treatment efficacy monitoring by bioluminescence of different groups of tumor-bearing mice.

### Oral ABC-101 reduces tumour proliferation and aggressiveness markers in *in vivo* human gastric cancer model

The antitumour activity of ABC-101 was evaluated in an orthotopic MKN-45_Luc human gastric cancer model established in Tg-hCD89 transgenic mice. Treatment with oral ABC-101 or vehicle was initiated two weeks after tumour-cell implantation into the stomach wall and continued for two weeks. At study endpoint, stomach tumours were analysed by immunohistochemistry for CD44, a marker associated with tumour migration and aggressiveness, and Ki67, a marker of cell proliferation (Suppl data: Figure 1). Oral ABC-101 significantly reduced CD44 expression, with a median staining score of 2.0±0.8 versus 3.5±0.6 in untreated animals (p=0.001). Notably, none of the seven ABC-101-treated animals reached the highest CD44 scores (3.5–4), compared with 8 of 12 untreated animals (Suppl data: Figure 1b.). Ki67 expression was also significantly reduced following ABC-101 treatment (3.0±0.5 vs 3.9±0.3; *p*=0.009) (Suppl data: Figure 1c.). Strikingly, tumour cells at the periphery of several ABC-101-treated lesions showed complete loss of CD44 or Ki67 staining (Suppl data: Figure 2, black arrows), a pattern not observed in untreated tumours. Together, these findings show that oral ABC-101 treatment is associated with reduced expression of markers of tumour aggressiveness and proliferation in this gastric cancer model.

### Oral ABC-101 Plus Irinotecan achieves superior antitumor activity compared with Cetuximab/FOLFIRI, outperforming current standard of care

The therapeutic potential of ABC-101 in combination with chemotherapy was next evaluated using treatment regimens representative of first-line therapy for metastatic colorectal cancer. Mice received irinotecan (200 mg/kg, intraperitoneally) or FOLFIRI regimen, consisting of irinotecan (200 mg/kg) plus 5-fluorouracil (40 mg/kg). Chemotherapy was administered either alone or in combination with intravenous cetuximab (4 mg during the first week followed by 2.5 mg once weekly for the subsequent 2 weeks) or with orally administered ABC-101 (0.2 or 0.8 mg/day), given once daily by gavage, 5 days per week for 3 weeks.

Among all treatment groups, combination of ABC-101 (0.8 mg/day) and irinotecan induced highest inhibition of primary tumor growth, yielding the lowest mean cecal tumor weight (0.095 ± 0.003 g), corresponding to a 76% reduction compared to untreated animals (0.404 ± 0.212 g; Figure 6b.). Adding ABC-101 significantly enhanced the therapeutic efficacy of irinotecan given alone, reducing tumor weight from 0.148 ± 0.001 g to 0.095 ± 0.003 g (p < 0.0001). Similarly, combining ABC-101 with FOLFIRI further reduced tumor weight compared with FOLFIRI monotherapy (0.095 ± 0.003 g *vs*. 0.126 ± 0.010 g; p = 0.019). Importantly, the ABC-101/irinotecan combination also outperformed the standard of care cetuximab/FOLFIRI regimen (0.095 ± 0.003 g *vs*. 0.121 ± 0.004 g; p= 0.0027), representing an additional reduction of approximately 22% in primary tumor burden. Collectively, these findings establish oral ABC-101 plus irinotecan as the most effective therapeutic combination tested.

### Oral ABC-101 potently suppresses distant pulmonary metastases and enhances chemotherapy efficacy

Pulmonary metastatic burden was quantified by flow cytometric analysis of human CD44-positive HT-29 cells isolated from lung tissue (Figure 6c.). In untreated animals, CD44-positive tumor cells represented 0.470 ± 0.054% of total lung cells. Oral administration of ABC-101 (0.8 mg/day) reduced this proportion to 0.123 ± 0.014%, corresponding to a 74% reduction in pulmonary metastatic burden compared with untreated controls (p < 0.0001). The therapeutic benefit of ABC-101 was further evaluated in combination with irinotecan or FOLFIRI. The addition of ABC-101 to irinotecan reduced the proportion of pulmonary CD44-positive tumor cells by 68% compared with irinotecan alone (p < 0.0001). Likewise, combining ABC-101 with FOLFIRI reduced pulmonary tumor burden by 59% relative to FOLFIRI monotherapy (0.138 ± 0.031% *vs*. 0.211 ± 0.059%; p = 0.0003). Compared with untreated animals, the combinations of ABC-101 with irinotecan or FOLFIRI reduced pulmonary metastatic colonization by approximately 5.5-fold and 3.4-fold, respectively. Among all combination regimens, ABC-101 plus FOLFIRI yielded the lowest mean pulmonary tumor cell burden (0.086 ± 0.015% CD44-positive cells), compared with 0.138 ± 0.080% in the ABC-101/irinotecan group. Collectively, these findings demonstrate that oral ABC-101 markedly limits pulmonary metastatic dissemination both as monotherapy and in combination with standard chemotherapy.

### Longitudinal monitoring of tumor progression by Bioluminescence Imaging

Therapeutic responses were further monitored by serial bioluminescence imaging throughout the treatment period (Figure 6d.). Oral administration of ABC-101 at 0.8 mg/day maintained the primary tumor bioluminescence signal close to baseline throughout the treatment period, indicating effective control of tumor progression. A similar, although less pronounced, effect was observed with the 0.2 mg/day dose. Combination therapy produced substantially greater antitumor activity: mice receiving ABC-101 plus irinotecan exhibited complete disappearance of the macroscopic bioluminescence signal from the primary tumor after the second week of treatment. In contrast, residual tumor signals remained detectable in animals treated with irinotecan or FOLFIRI alone. Similarly, mice treated with ABC-101 plus FOLFIRI displayed only minimal residual bioluminescence at the primary tumor site after 3 weeks of treatment, whereas persistent tumor signals remained evident in animals receiving the reference cetuximab/FOLFIRI regimen. At the end of the study, the combination of oral ABC-101 with irinotecan achieved the most effective control of disease progression, with no detectable macroscopic bioluminescence originating from either primary tumors or metastatic lesions.

### Oral ABC-101 drives robust recruitment of CD89⁺ myeloid effector cells to primary tumors and metastases

To investigate the cellular mechanisms underlying ABC-101 antitumor activity, CD89⁺ immune populations were analyzed by flow cytometry in primary cecal tumors (Figure 7, a.-d.) and liver and lung metastases (Figure 7, e.-h.). Mice receiving Irinotecan alone were compared with those treated with Irinotecan plus oral ABC-101 (0.8 mg/day).

**Figure 7:**
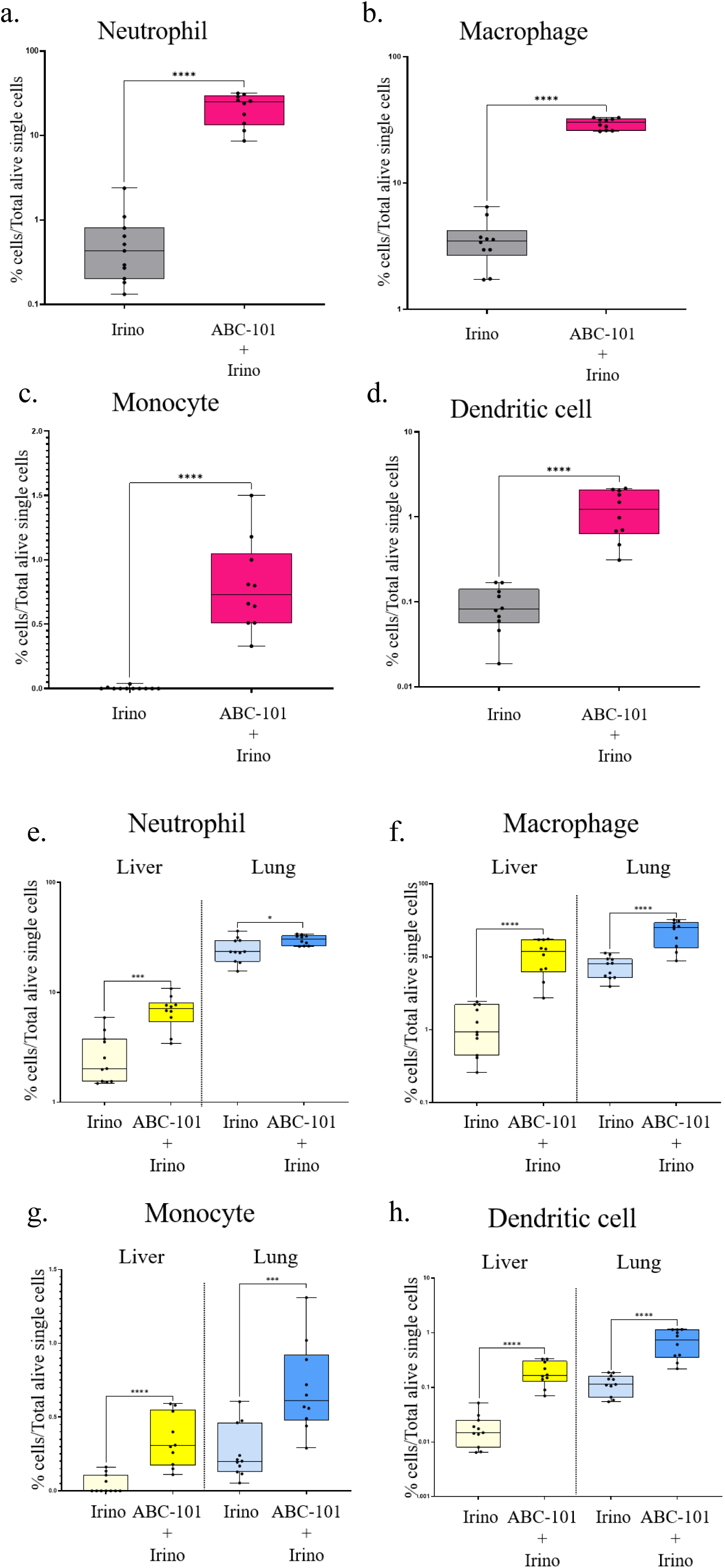
Recruitment of CD89^positive^ effector cells in tumoral foci. a – d: Recruitment of a. neutrophils, b. macrophages, c. monocytes and d. dentritic cells in primary tumor site of cecum; e-h: Recruitment of e. neutrophils, f. macrophages, g. monocytes and h. dentritic cells in Liver and Lung metastasis

The addition of oral ABC-101 to irinotecan resulted in a marked increase in the infiltration of CD89⁺ myeloid effector cells within primary tumors (Figure 7, a.-d.). The proportion of CD89⁺ Ly6G^high^ neutrophils increased from 0.63 ± 0.65% following irinotecan monotherapy to 21.87 ± 8.33% after combination treatment (p < 0.0001). Similarly, CD89⁺ Ly6G⁻ CD11b⁺ F4/80⁻ monocytes increased from 0.004 ± 0.010% to 0.79 ± 0.35% (p < 0.0001). The most pronounced change was observed in the macrophage population. CD89⁺ CD11c⁺ Ly6G⁻ F4/80⁺ macrophages represented 29.49 ± 2.97% of total tumor cells in mice receiving ABC-101 plus irinotecan, compared with only 3.57 ± 1.49% following irinotecan alone (p < 0.0001). Likewise, CD89⁺ CD11c^high^ dendritic cells increased from 0.09 ± 0.05% to 1.27 ± 0.73% (p < 0.0001).

A similar pattern was observed in metastatic lesions (Figure 7, e.-h.). In the liver, the proportion of CD89⁺ Ly6G^high^ neutrophils increased nearly nine-fold, from 1.24 ± 0.80% with irinotecan alone to 10.92 ± 5.50% following combination therapy (p < 0.0001). CD89⁺ monocytes also increased significantly, from 0.04 ± 0.06% to 0.34 ± 0.18% (p < 0.0001). Macrophage and dendritic cell populations exhibited comparable increases, with mean frequencies approximately 2.5-fold and 10-fold higher, respectively, than those observed after irinotecan monotherapy. In metastatic lungs, combination therapy similarly enhanced the recruitment of CD89⁺ myeloid cells. CD89⁺ Ly6G^high^ neutrophils increased from 7.50 ± 2.49% to 22.10 ± 8.38% of total lung cells. CD89⁺ monocytes increased from 0.26 ± 0.17% to 0.69 ± 0.31% (p = 0.0006), while CD89⁺ alveolar macrophages increased from 24.87 ± 6.20% to 30.12 ± 3.16% (p = 0.036). Similar increases were also observed in the CD89⁺ dendritic cell population.

In the caecum, oral ABC-101 combined with irinotecan significantly increased the recruitment of all four CD89-positive immune cell populations—neutrophils, macrophages, monocytes and dendritic cells—compared with cetuximab, either as monotherapy or combined with FOLFIRI (Suppl. data: Figure 3a.-d.). A similar trend was observed in liver and lung metastases, with greater neutrophil recruitment following ABC-101-based combination therapy than with cetuximab (Suppl. data: Figure 3e.).Together, these results show that oral ABC-101 profoundly reshapes the tumor immune microenvironment by recruiting CD89⁺ myeloid effector cells at both primary and distant metastatic sites.

## DISCUSSION

ABC-101 demonstrated high specificity for CEA-overexpressing tumor cells across the four gastrointestinal cancers evaluated, supporting CEA as a clinically relevant therapeutic target. The highest expression was observed in colorectal cancer, particularly in rectal tumors and stage III–IV colon cancers. Importantly, CEA expression increased with disease progression, with higher median ABC-101 labeling in stage IV versus stage III tumors, suggesting greater target availability in metastatic disease. CEA has also been reported to remain stable in metastatic lesions despite prior chemotherapy or radiotherapy (20,21), reinforcing its value throughout disease progression. Although expression was more heterogeneous in esophageal and gastric cancers, more than half of tumor cells expressed CEA, identifying additional populations potentially eligible for ABC-101.

Overall, the correlation between CEA expression and ABC-101 labeling confirms target specificity, while broad expression across malignant cells may reduce antigen-negative escape. These findings provide a strong rationale for evaluating ABC-101 particularly in advanced colorectal cancer. This study provides the first comprehensive characterization of the pharmacokinetics and biodistribution of orally administered secretory immunoglobulin A (SIgA) in healthy and tumor-bearing mice. Gastrointestinal transit in mice is normally completed within approximately 6 hours, with clearance of luminal contents about 2 hours later (15). In contrast, ABC-101 reached a plasma Tmax of approximately 4 hours and remained detectable in gastrointestinal mucosa and several peripheral organs for up to 24 hours, indicating resistance to the rapid proteolytic degradation typically associated with dietary proteins.

The rapid appearance of ABC-101 in circulation strongly suggests active intestinal uptake. SIgA is known to cross the intestinal epithelium through receptor-mediated mechanisms (4) and intestinal loop experiments detected ABC-101 in blood within approximately one hour of luminal administration. Although circulating ABC-101 rapidly became undetectable after its peak, the antibody persisted in tissues for up to 24 hours. This differs markedly from conventional IgG antibodies, which persist in the vascular compartment but show comparatively limited tissue penetration. Lacking FcRn interaction (16), SIgA appears to undergo rapid systemic distribution followed by tissue localization rather than prolonged circulation. This profile may provide safety advantages over intravenously administered IgG antibodies by limiting prolonged systemic exposure while favoring tissue distribution. Although this remains to be confirmed in dedicated toxicology studies, the present findings support a potentially favorable pharmacokinetic profile for orally administered ABC-101.

Biodistribution was markedly different in healthy and tumor-bearing animals. In healthy mice, ABC-101 was initially concentrated in the gastrointestinal tract and rapidly cleared from digestive and systemic tissues. In tumor-bearing mice, however, ABC-101 reached the primary cecal tumor with a Tmax of approximately 4 hours and remained detectable for at least 96 hours. ABC-101 also accumulated at distant metastatic sites, with concentrations at 24 hours approximately 10-fold higher in metastatic liver and 20-fold higher in metastatic lungs than in corresponding healthy organs, together with prolonged retention in mesenteric lymph nodes and peritoneal lesions. Thus, ABC-101 is not restricted to the gastrointestinal tract after oral administration but selectively accumulates and persists in CEA-positive primary tumors and distant metastases. This pattern supports target-dependent tumor retention and provides a mechanistic basis for sustained antitumor activity despite short systemic exposure.

The ABC-101 dose was selected in part by reference to the cetuximab dosing regimen. The recommended human cetuximab regimen consists of a 400 mg/m² induction dose followed by 250 mg/m² weekly maintenance. Based on an estimated mouse absorption surface area of 0.01 m², this corresponds to 4 mg during the first week followed by 2.5 mg/week. Because SIgA was administered orally for five consecutive days per week, 0.8 mg/day was selected, corresponding to one-fifth of the 4 mg induction dose. A four-fold lower dose (0.2 mg/day) was also evaluated to assess dose response. Orally administered ABC-101 demonstrated robust antitumor activity both as monotherapy and in combination with standard chemotherapy. In the orthotopic metastatic colorectal cancer model, ABC-101 significantly reduced primary cecal tumor burden and pulmonary metastases, outperforming both the corresponding anti-CEA IgG1 and cetuximab. Therapeutic activity was further enhanced when ABC-101 was combined with irinotecan or FOLFIRI. Compared with chemotherapy alone, combination treatment consistently reduced primary tumor burden, pulmonary metastases and overall bioluminescent tumor burden. ABC-101 plus irinotecan produced the strongest activity across efficacy endpoints. Importantly, these effects extended to distant metastatic sites, consistent with prolonged ABC-101 retention within CEACAM5-positive lesions. Although the study was not designed to formally establish pharmacological synergy, the consistent improvement across independent efficacy endpoints supports the therapeutic benefit of combining ABC-101 with irinotecan and provides a rationale for clinical evaluation in metastatic colorectal cancer, including comparison with current irinotecan-based regimens.

The efficacy studies were performed in SCID mice lacking functional T and B lymphocytes, indicating that ABC-101 activity relies predominantly on innate immune mechanisms involving FcαRI (CD89)-expressing myeloid cells. Compared with irinotecan alone, addition of ABC-101 consistently increased CD89-positive neutrophils, monocytes, macrophages and dendritic cells in primary tumors and hepatic and pulmonary metastases. The strongest effect involved neutrophils, with an approximately 35-fold increase in primary tumors and 3-to 10-fold increases at metastatic sites. This enhanced myeloid infiltration was associated with greater control of tumor growth and metastatic dissemination, although causality was not directly established. The observation is consistent with FcαRI biology in humans and its recapitulation in the hFcαRI (hCD89) transgenic mouse model (17). Engagement of CD89 by IgA activates neutrophils, monocytes, macrophages, and dendritic cells, triggering ADCC, ADCP, trogoptosis, and inflammatory mediator release (18,19). Activated neutrophils can further amplify this response through cytokines and chemokines, promoting additional myeloid recruitment and activation, even in SCID genetic background (20). Importantly, increased CD89-positive effector-cell infiltration occurred in both primary tumors and liver and lung metastases, where ABC-101 also showed prolonged retention. This spatial association supports target-dependent activation of innate immunity at both primary and metastatic sites. Remarkably, substantial antitumor activity was achieved in the absence of adaptive immunity, demonstrating that oral SIgA can mobilize innate mechanisms alone to control primary and metastatic tumors.

These findings support orally administered SIgA as a therapeutic platform with pharmacological and biological properties fundamentally different from intravenous IgG. ABC-101 combines selective retention within CEACAM5-positive tumors and metastases, recruitment of FcαRI-expressing myeloid effector cells and superior antitumor activity compared with the IgG1 antibodies evaluated here. This concept is supported by evidence that immune events initiated within the intestinal mucosa can influence systemic antitumor immunity. Zitvogel and colleagues established the role of the intestinal microbiota and mucosal immune system in responses to anticancer therapies (21)), while Devaud and colleagues showed that immune activation within intestinal tumors may be associated with immune responses in distant liver metastases (22). Although neither study investigated therapeutic secretory IgA, both support the concept that immune activation initiated in the intestinal mucosa is not confined to the gut but may influence distant metastatic sites. Similarly, the pioneering work of van Egmond (23–26), Valerius (27–32), Leusen (30,33–35) and others established FcαRI (CD89) as one of the most potent activating Fc receptors on human myeloid cells and demonstrated the capacity of multimeric IgA immune complexes to activate these populations. The increased infiltration of CD89-positive cells observed here is therefore compatible with a major contribution of FcαRI-dependent innate immunity to ABC-101 activity, although a direct mechanistic relationship remains to be established.

Human immunology also suggests that SIgA may engage pathways beyond innate myeloid immunity. SIgA is a physiological ligand of FCRL3 on subsets of regulatory T cells, where it may modulate their suppressive phenotype. Moreover, pIgR-expressing ovarian cancer cells can locally generate SIgA from tumor-specific dimeric IgA, potentially influencing the tumor immune microenvironment (8,36).These mechanisms could not be addressed in the SCID model but suggest that adaptive immunity may further amplify SIgA activity in immunocompetent hosts. Studies in immunocompetent or humanized models will be required to define these contributions.

The biological activity of ABC-101 may not be restricted to colorectal cancer. Preliminary results in a second human tumour model using CEACAM5^high^ MKN-45 gastric cancer cells showed that oral ABC-101 significantly reduced CD44 and Ki67 expression, markers associated with tumour aggressiveness/migration and proliferation, respectively. Some tumour cells completely lost CD44 or Ki67 staining, suggesting marked changes in their proliferative and aggressive phenotype. Although the underlying mechanisms remain to be determined, these findings indicate that ABC-101 can influence tumour-cell biological programmes beyond its effects on tumour burden and immune-cell recruitment. This observation in an independent gastrointestinal cancer model reinforces the antitumour activity demonstrated in metastatic colorectal cancer and supports the broader potential of ABC-101 in CEACAM5-expressing gastrointestinal malignancies. Gastric cancer therefore represents a relevant indication for further investigation, particularly from the second-line setting, where therapeutic options remain limited, and unmet medical need is critical.

In conclusion, these findings identify orally administered SIgA as a promising immunotherapeutic platform combining targeted delivery, tumor-selective tissue retention, and activation of FcαRI-dependent innate immunity. Beyond ABC-101, they support the broader concept that the intestinal mucosa can serve as a therapeutic gateway to control both gastrointestinal primary tumors and distant metastatic disease. If confirmed in future preclinical and clinical studies, oral SIgA could establish a new class of antibody-based cancer immunotherapy.

## MATERIAL and METHODS

### Animals

All animal experiments were conducted in accordance with institutional guidelines and national regulations governing the care and use of laboratory animals. Experimental protocols were approved by the appropriate institutional animal ethics committee and performed in compliance with applicable legislation. SCID-CD89 transgenic mice (10-16 weeks old) were used in collaboration with CRIBL_CNRS 7276_INSERM 1262 laboratory from Limoges University.

### Reagents

Clinical-grade cetuximab (Erbitux®, Merck), irinotecan, and 5-fluorouracil (5-FU) were obtained from the Pharmacy Department of Limoges University Hospital and used according to the manufacturers’ recommendations.

ABC-101, a recombinant secretory human IgA directed against CEACAM5, was produced by Icosagen (Estonia). Production batches #1 and #2 throughout the study.

### Flow cytometry antibodies

The following fluorochrome-conjugated monoclonal antibodies were used for flow cytometry analyses:

• anti-CD11c-BV421 (Biolegend, clone N418)

• anti-CD44-PE (Biolegend, clone BJ18)

• anti-F4/80-PE-Dazzle 594 (Biolegend, clone BM8)

• anti-NKp46-AF700 (BD Biosciences, clone 29A1.4)

• anti-CD11b-BV605 (Biolegend, clone M1/70)

• anti-Ly6G-FITC (BD Biosciences, clone 1A8)

• anti-CD63-PC7 (Biolegend, clone NVG-2)

• anti-human IgA-AF647 (SouthernBiotech)

### Generation of anti-CEACAM5 monoclonal antibodies

HAMIGA™ mice (6–8 weeks of age) were immunized subcutaneously with 10 µg of purified human carcinoembryonic antigen (hCEACAM5; Abcys, France) emulsified in Freund’s Complete Adjuvant. Booster immunizations were administered every two weeks using Freund’s Incomplete Adjuvant. A final booster injection was performed four days before spleen collection. Splenocytes were isolated under sterile conditions and fused with SP2/0 myeloma cells using polyethylene glycol, according to the method originally described by Köhler and Milstein. Hybridomas were cultured in high-glucose DMEM supplemented with fetal calf serum, hybridoma medium, OPI supplement, recombinant murine IL-6 and hypoxanthine/thymidine. Hybridoma selection was performed using HAT medium containing aminopterin for two weeks. Surviving hybridomas were expanded in complete DMEM supplemented with fetal calf serum, penicillin, streptomycin and recombinant murine IL-6. Culture supernatants were screened for reactivity against recombinant human CEACAM5 by ELISA. Positive hybridomas were subjected to limiting dilution of cloning to ensure monoclonality. Stable antibody-producing clones were subsequently expanded and characterized for antigen specificity, antibody production, and growth characteristics.

### Anti-CEA ELISA

High-binding 96-well ELISA plates (MaxiSorp®, Thermo Fisher Scientific) were coated overnight at 4°C with recombinant human CEACAM5 (1 µg/mL) diluted in phosphate-buffered saline (PBS). After washing, nonspecific binding sites were blocked with 3% bovine serum albumin (BSA) in PBS. Hybridoma culture supernatants or purified ABC-101 were added to the wells and incubated for 1 hour at 37°C. Following washing, bound antibodies were detected using horseradish peroxidase- or alkaline phosphatase-conjugated anti-human IgA secondary antibodies. After substrate enzymatic cleavage, absorbance was measured at 450 nm. Samples were considered positive when the absorbance exceeded the predefined cutoff established from negative-control wells.

### SDS-PAGE analysis

The structural integrity of ABC-101 was assessed by SDS-PAGE using NuPAGE™ 3–8% Tris-Acetate gels (Invitrogen) according to the manufacturer’s instructions. Proteins were visualized using Colloidal Blue staining, and molecular weights were estimated using HiMark™ prestained protein standards (Invitrogen).

### Radiolabeling and stability studies

The physicochemical stability of ^125^I-labeled ABC-101 was evaluated under storage and physiological conditions. Storage stability of both the radiolabeling solution and the final dosing formulation (0.1 M glycine, 150 mM NaCl, pH 3.5) was assessed after storage at 4°C and at room temperature for up to 8 days. The stability of radiolabeled ABC-101 under physiological conditions was evaluated by incubating the antibody in freshly collected mouse blood at 37°C. Radiochemical purity was monitored over time using instant thin-layer chromatography (iTLC) and size-exclusion high-performance liquid chromatography (SE-HPLC).

To evaluate the effect of pH on antibody stability, the radiolabeling solution was diluted 1:10 into acidic (0.1 M glycine, 150 mM NaCl, pH 2.0), neutral (0.1 M phosphate buffer, pH 7.4), or basic (0.1 M Tris-HCl, pH 9.0) buffers and incubated at room temperature for 4 hours. Radiochemical integrity was assessed by iTLC and SE-HPLC.

### Pharmacokinetic and biodistribution studies

Healthy and tumor-bearing mice were randomly assigned to four experimental groups corresponding to sampling times of 4, 24, 48 and 96 hours after administration. To prevent uptake of free radioactive iodine by the thyroid and gastric mucosa, mice received potassium iodide (1 mg/kg) by oral gavage 24 hours and 1 hour before administration of ^125^I-ABC-101. Radiolabeled ABC-101 was equilibrated to room temperature before administration and delivered by oral gavage using flexible feeding tubes. At each scheduled time point, approximately 100 μL of blood was collected from the retro-orbital sinus under isoflurane anesthesia, and radioactivity was quantified using an automated gamma counter (Hidex AMG). At the terminal time point, mice were euthanized by intraperitoneal administration of ketamine (200 mg/kg) and xylazine (20 mg/kg). Peritoneal lavage was performed by injecting 5 mL of ice-cold high-glucose DMEM into the peritoneal cavity, followed by gentle abdominal massage. Lavage fluid was collected for radioactivity measurements. Organs of interest, including liver, lungs, small intestine, colon, cecal tumor, mesenteric lymph nodes, peritoneal tumor deposits and brain, were harvested, rinsed with physiological saline, blotted dry and weighed individually. Tissue-associated radioactivity was quantified using a calibrated automated gamma counter (Hidex AMG). Results were expressed as tissue radioactivity normalized to organ weight.

### Immunohistochemistry

Formalin-fixed, paraffin-embedded human tumor specimens were sectioned at a thickness of 4 μm and mounted onto positively charged glass slides. Following deparaffinization and rehydration, endogenous peroxidase activity was quenched with 3% hydrogen peroxide. After blocking nonspecific binding, tissue sections were stained using an anti-CEACAM5 monoclonal antibody corresponding to the IgG version of ABC-101 (Leica diagnostic antibody) on a Leica Bond RX automated immunostainer according to the manufacturer’s protocol. CEA expressions were evaluated by light microscopy. Both staining intensity and the proportion of positive tumor cells were independently assessed to determine overall CEACAM5 expression within each specimen.

### *In vivo* ileal loop assay

To evaluate intestinal uptake of ABC-101 *in vivo*, mice were fasted overnight before surgery. Animals were anesthetized with ketamine (100 mg/kg) and xylazine (10 mg/kg) and maintained at 37°C throughout the procedure. A 1.5-cm ileal loop containing a Peyer’s patch was isolated, and 100 µL of ABC-101 (1 mg/mL in PBS) was injected into the intestinal lumen. Control animals received only PBS vehicles (1 mg/mL). After 90 minutes, mice were euthanized, and the intestinal segment was extensively washed with PBS before overnight fixation in 4.5% paraformaldehyde. Tissues were embedded in OCT compound, cryosectioned (7 µm), and mounted onto Superfrost Ultra Plus microscope slides. ABC-101 localization was visualized by immunofluorescence using a fluorochrome-conjugated anti-human IgA antibody. Images were acquired using a Nikon Eclipse Ti2 fluorescence microscope (magnitude x10 objective).

### Orthotopic colorectal cancer model

An orthotopic colorectal cancer model was developed using HT29-luc human colorectal adenocarcinoma cells. Briefly, mice aged 2.5–4 months were anesthetized with inhaled isoflurane, and a midline laparotomy was performed under sterile conditions. The cecum was gently exteriorized, and 1×10^5^ HT29-luc cells suspended in 10 µL PBS were injected into the cecal wall under stereomicroscopic guidance using a 30-gauge insulin syringe. Following injection, the cecum was returned to the abdominal cavity, and the abdominal wall was closed in anatomical layers. This orthotopic model reproduces local tumor growth together with spontaneous metastatic dissemination, thereby closely mimicking the natural progression of human colorectal cancer.

### Orthotopic gastric cancer model

Mice were anesthetized with isoflurane and positioned in the right lateral decubitus position to expose the stomach. MKN45 diffuse gastric cancer cells stably expressing firefly luciferase (MKN45-LucF) were orthotopically injected into the gastric wall at a dose of 10,000 cells suspended in 30 μL of Matrigel (7 mg/mL). Following cell implantation, the muscular layer and skin were closed using standard suturing procedures.

### Immunohistochemistry and Histopathological Analysis

Serial 3-μm-thick sections were prepared from formalin-fixed, paraffin-embedded stomach tissues. Immunohistochemical staining was performed using antibodies directed against human CD44 and Ki67 according to previously validated protocols (37).

Immunostaining was evaluated using a semi-quantitative scoring system that assessed both the intensity and the extent of staining across the entire tissue section. Slides were independently scored in a blinded manner by Coralie Genevois and Christine Varon according to the following criteria:

• Score 1 (Weak): predominantly weak staining in the majority of tumor cells.

• Score 2 (Moderate): heterogeneous staining, with approximately 50% weakly stained cells and 50% strongly stained cells.

• Score 3 (Extensive): heterogeneous staining, with ≥50% of tumor cells showing strong staining.

• Score 4 (Severe): homogeneous strong staining in >90% of tumor cells.

### Treatment in human gastric cancer model

Oral treatment with ABC-101 begins 15 days after transplantation. The treatment regimen consists of two cycles of five oral administrations of 0.8mg (via gavage in awake mice) per week for two weeks, with two treatment-free days at the weekend. The animals are euthanised three days after the final oral administration. The stomach is then removed, and the organ is embedded in paraffin for immunohistochemical analysis.

### Bioluminescence imaging

Tumor progression was monitored longitudinally by bioluminescence imaging. Mice received an intraperitoneal injection of D-luciferin (2 mg in 200 µL) and were anesthetized with isoflurane. Bioluminescence images were acquired using an *in vivo* imaging system beginning 11 minutes after substrate administration. Total photon flux (photons/s) was quantified using the manufacturer’s analysis software. Tumor burden was monitored weekly by both bioluminescence imaging and caliper measurements.

### Flow cytometry

Single-cell suspensions were prepared from primary cecal tumors, liver, and lung tissues by mechanical dissociation followed by enzymatic digestion with collagenase. Cell suspensions were filtered through 40-µm cell strainers, and erythrocytes were lysed. Cells were subsequently washed and resuspended in flow cytometry staining buffer. After cell counting, nonspecific Fc receptor binding was blocked using an Fc receptor–blocking reagent. Cells were then stained with a viability dye together with fluorochrome-conjugated monoclonal antibodies specific for myeloid lineage markers. Flow cytometric analyses were performed on viable singlet cells following sequential exclusion of debris, doublets, and dead cells. Neutrophils were identified as CD11b^+^Ly6G^+^ cells, inflammatory monocytes as CD11b^+^Ly6G^-^F4/80^-^ cells, macrophages as CD11b^+^F4/80^+^Ly6G^-^ cells, and dendritic cells as CD11c^high^Ly6G^-^F4/80^-^ cells. Fluorescence-minus-one (FMO) controls, together with unstained and single-stained compensation controls, were used to establish gating strategies and fluorescence compensation. Data were acquired using a CytoFLEX (Beckman Coulter) and analyzed with BD FACSDiva™ Software (BD Biosciences). Cell populations were expressed as the percentage of viable leukocytes recovered from each organ.

### Statistical analysis

Data are presented as mean ± standard deviation (SD) unless otherwise indicated. Statistical analyses were performed using GraphPad Prism 10 (GraphPad Software, San Diego, CA, USA). Comparisons between two independent groups were performed using the two-tailed Mann–Whitney *U* test. All tests were two-sided, and *P* values <0.05 were considered statistically significant.

Statistical significance is indicated as follows:

- ns, not significant
- *\* p* < 0.05
- *\*\* p* < 0.01
- *\*\*\* p* < 0.001
- *\*\*\*\* p* < 0.0001

### Study design

Animals were randomly allocated to different experimental groups. Investigators performing histological, imaging, and flow cytometric analyses were blinded to treatment allocation whenever feasible. No animals were excluded from the analyses unless predefined experimental criteria were met. Sample sizes were determined based on previous experience with the orthotopic colorectal cancer model and were sufficient to detect biologically meaningful differences between treatment groups.

## Acknowledgements

We would like to thank Dr Eric Pinaud for making the research facilities at the CRIBL laboratory available to us. We are grateful to BISCEm unit (Univ. Limoges, UAR 2015 CNRS, US 42 Inserm, CHU Limoges) for technical support regarding microscopy, histology and cytometry experiments and animal core facility.

## Contributors

Conceptualization: MB, JMH and AC Methodology: MB, SP, CV, AC. Formal Analysis: MB, JMH, AC. Investigation: MB, AP, OFB, BC, CG, AC. Resources: OFB, SP, CV, JMH,AC. Writing Original Draft: AC. Writing—Review and Editing: MB, SP, CV, BL, JMH, AC. Supervision and Funding Acquisition: AC, JMH. Guarantor: AC.

