## Supplemental data figure for "Beyond IgG: Oral Secretory IgA as a new efficient modality for cancer immunotherapy"

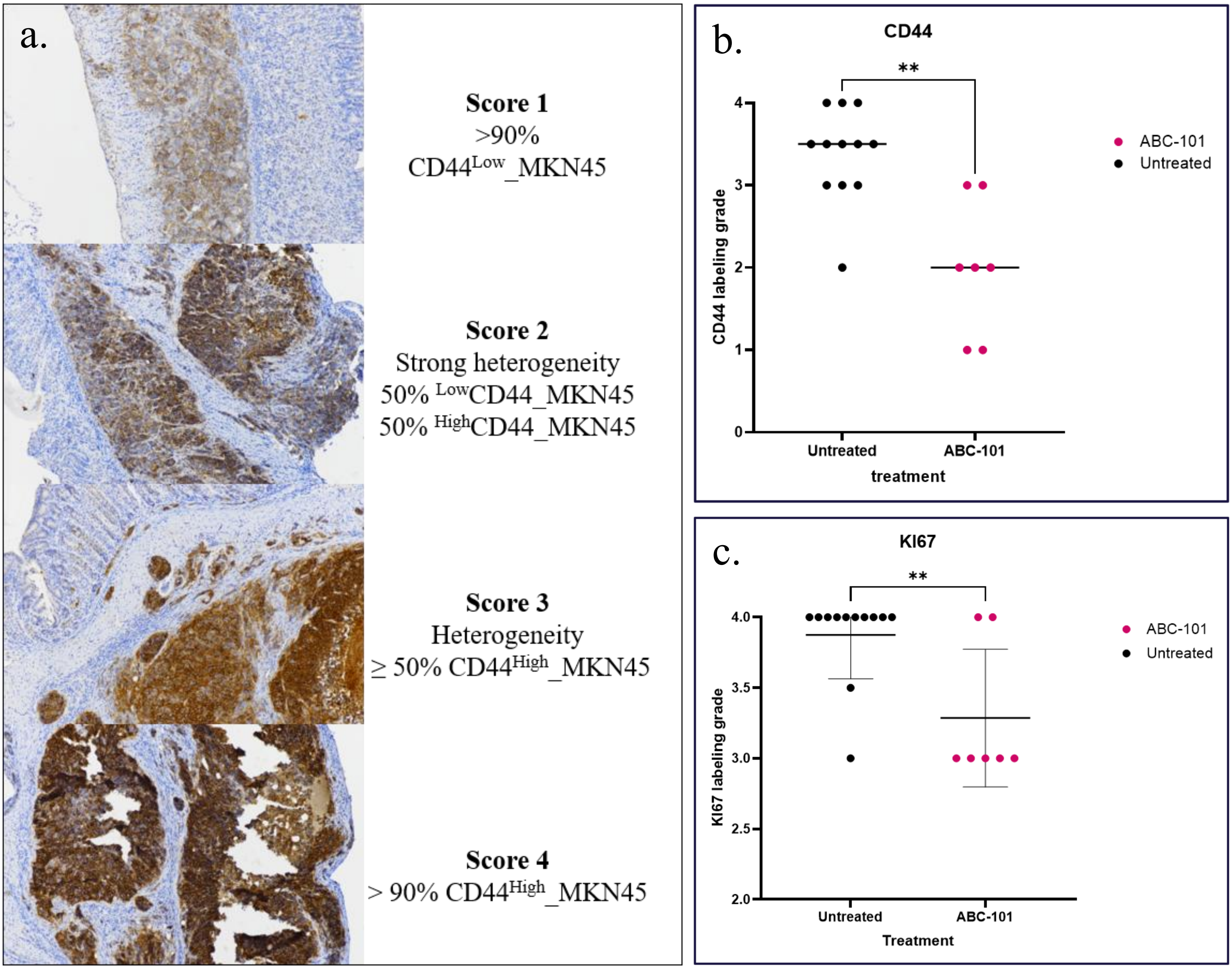


**Supplementary Data: Figure 1 CD44 and KI67 labeling scoring of CEA$^{High}$ MKN45 cells in tumoral gastric epithelium**

a. Relative quantification of CD44 expression in gastric cancer (GC) tissues b. Scoring of CD44 labeling in immunohistochemistry in GC tissues of oral ABC-101 and untreated groups. c. Scoring of KI67 labeling in immunohistochemistry in GC tissues of oral ABC-101 and untreated groups..

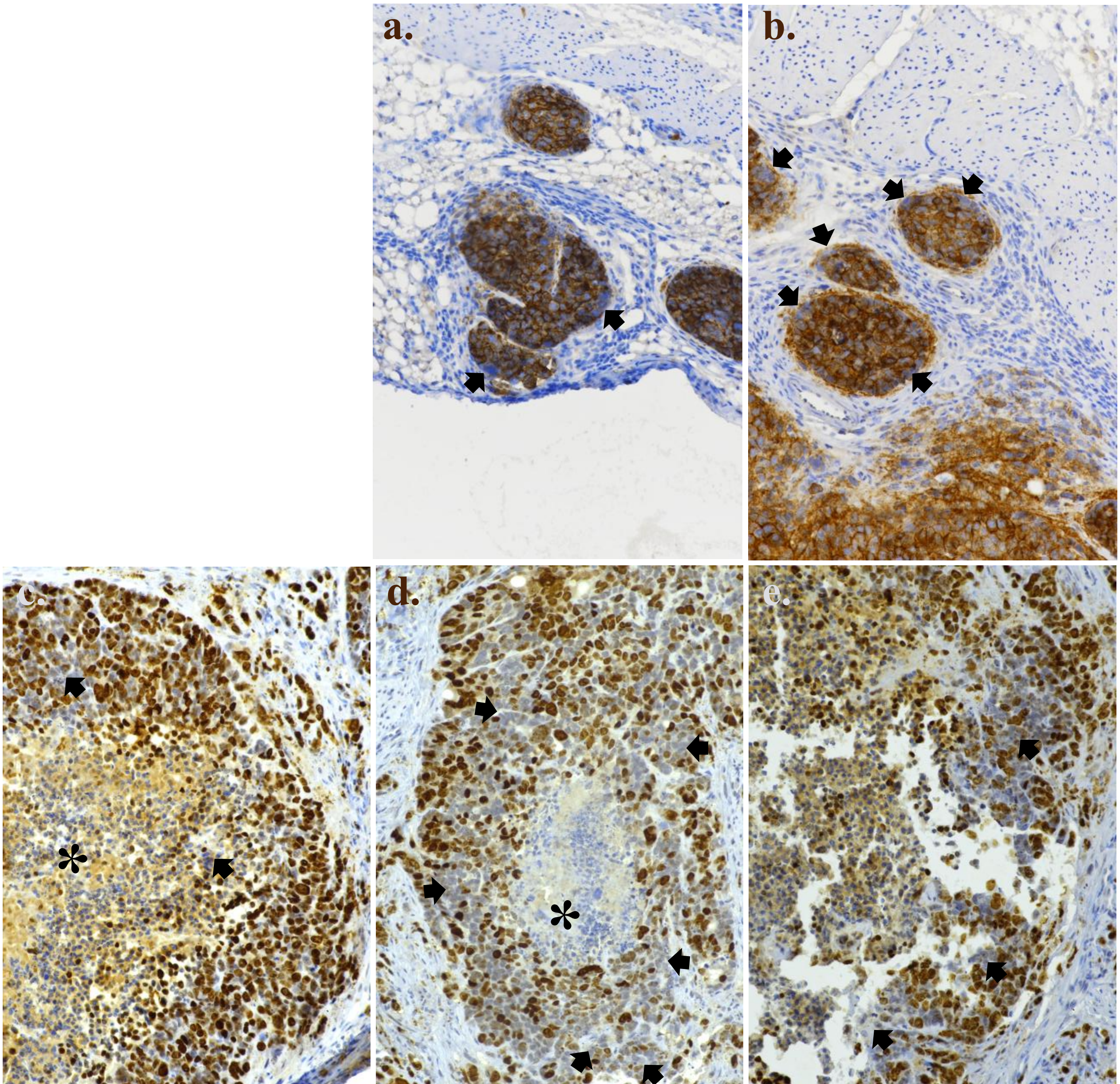


**Supplementary Data: Figure 2 CD44 and KI67 negatived CEA$^{High}$ MKN45 cells in tumoral gastric epithelium by oral ABC-101 treatment**.

A-B: Representative images of immunohistochemistry staining of CD44 in gastric cancer (GC) tissues of ABC-101 treated group. Black arrows focused on CD44$^{negative}$ MKN45 cells; C-E: Representative images of immunohistochemistry staining of KI67 in GC tissues of ABC-101 treated group. Black arrows focused on KI67$^{negative}$ MKN45 cells, black star focused on necrotic foci; F-H: Representative images of immunohistochemistry staining of KI67 in GC tissues of untreated group.

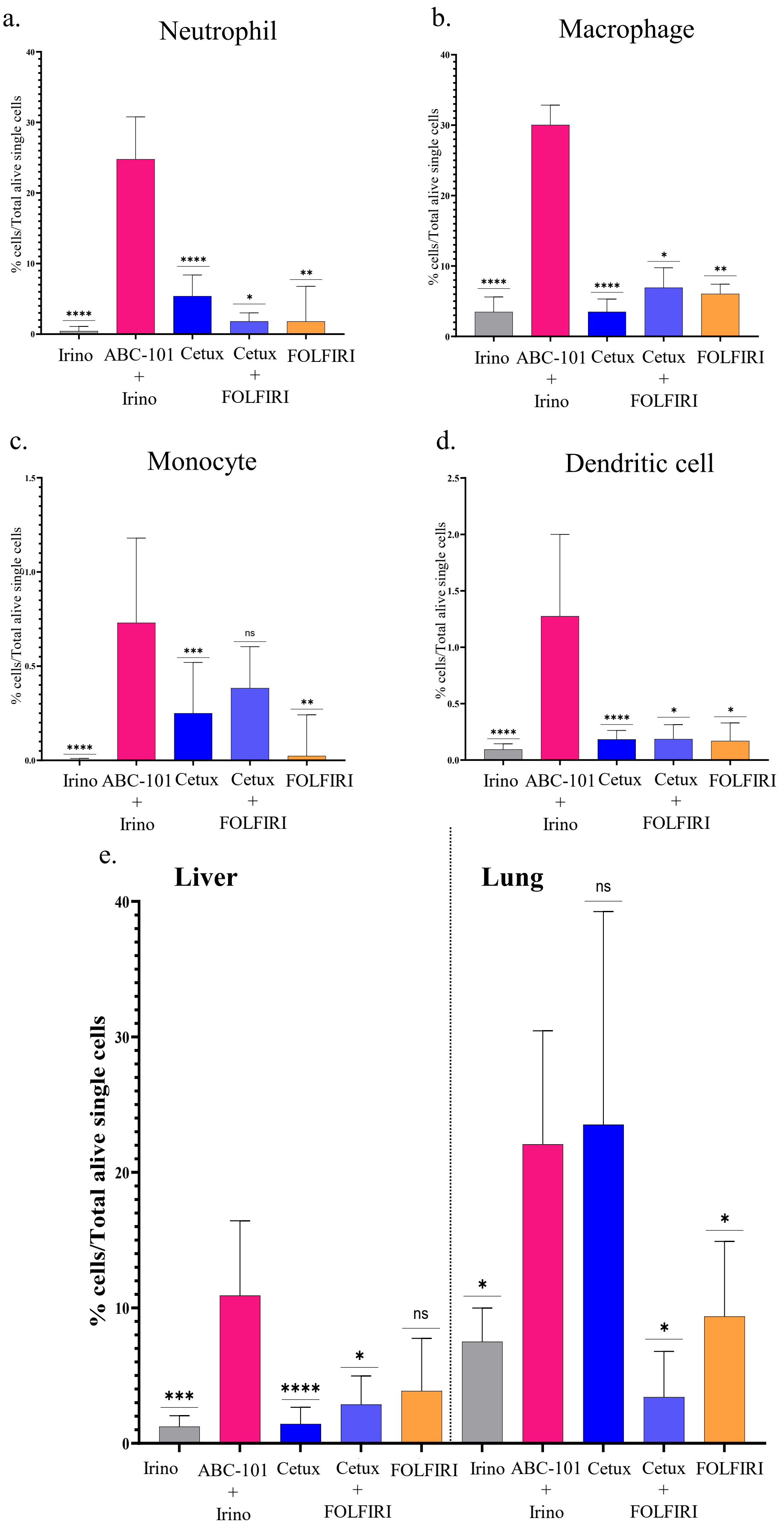


**Supplementary Data: Figure 3 Recruitment of CD89$^{positive}$ effector cells in tumoral foci in Irinotecan, ABC-101/Irinotecan, Cetuximab, Cetuximab/FOLFIRI and FOLFIRI treated groups**

a – d: Recruitment of a. neutrophils, b. macrophages, c. monocytes and d. dentritic cells in primary tumor site of cecum; e: Recruitment of neutrophils, in Liver and Lung metastasis. The statistical significance of each group (Irino, n=11; Cetux, n=9; Cetux+FOLFIRI, n=2; FOLFIRI, n=4) was assessed by comparison with the ABC-101/irinotecan-treated group (n=10).
